# A Machine-learning-based Method to Detect Degradation of Motor Control Stability with Applications to Diagnosis of Presymptomatic Parkinson’s Disease

**DOI:** 10.1101/2022.10.17.512460

**Authors:** Vrutangkumar V. Shah, Shail Jadav, Sachin Goyal, Harish J. Palanthandalam-Madapusi

**Affiliations:** Balance Disorder Lab, Department of Neurology, Oregon Heath and Science University, OR - 97239, USA; SysIDEA Robotics Lab, Mechanical Engineering, Indian Institute of Technology (IIT) Gandhinagar, GJ-382055, India; Department of Mechanical Engineering, University of California, Merced, CA-95343, USA; Health Science Research Institute, University of California, Merced, CA-95343, USA

## Abstract

Parkinson’s disease (PD), a neuro-degenerative disorder, is often detected by onset of its motor symptoms such as rest tremor. Unfortunately, motor symptoms appear only when approximately 40%-60% of the dopaminergic neurons in the substantia nigra are lost. In most cases, by the time PD is clinically diagnosed, the disease may already have started 4 to 6 years beforehand. So there is a need for developing a test for detecting PD *before* the onset of the motor symptoms. This phase of PD is referred to as Presymptomatic PD (PPD). The motor symptoms of Parkinsons Disease are manifestations of instability in the sensorimotor system that develops gradually due to the neuro-degenerative process. In this paper, based on the above insight, we propose a new method that can potentially be used to detect degradation of motor control stability which can be employed for the detection of PPD. The proposed method tracks the tendency of a feedback control system to transition to an unstable state, and uses machine learning algorithm for its robust detection. This method is explored using simulations of a simple pendulum with PID controller as a conceptual representation for both healthy and PPD individuals. We also propose an example task with physiological measurements that can be used with this method and potentially be employed in a clinical setting. We present representative data collected through such a task, thereby demonstrating the feasibility to generate data for the proposed method.

**Author summary:** Parkinson’s disease (PD) is a neuro-degenerative disorder that develops and progresses over several years. Currently, one is able to diagnose PD only after the appearance of motor symptoms (symptoms in movements of body parts), which unfortunately may be 4 to 6 years after the neuro-degeneration may have started. It has been shown that there are benefits to diagnosing PD at early stages, motivating the need to explore tools for diagnosing PD in the pre-symptomatic stage referred to as Presymptomatic Parkinson’s disease (PPD). In this paper, a novel approach is explored that utilises the insight that the motor symptoms in PD may be seen as an instability in the feedback-control system that controls movements of body parts (sensory-motor loop). The proposed method uses a series of simple movement tasks performed by an individual in a clinic as the input to detect any gradual degradation of movement control that is leading to an instability, but before the instability and consequently the symptoms are manifested. This method is tested through extensive simulations and a potential experimental realisation with preliminary data. While a full-fledged validation will be undertaken as part of future work, initial results show promise and feasibility of further data collection.

## 1 Introduction

Parkinson’s disease (PD) is the second largest progressive neurodegenerative disorder of the central nervous system [1]. It is characterized by several motor symptoms such as tremor at rest, rigidity, bradykinesia, and postural instability. Generally, the clinical diagnosis is based on a combination of clinical symptoms, a thorough history of patients, and response to levodopa [1–3]. It is reported that motor symptoms appear only when approximately 40%-60% of the dopaminergic neurons in the substantia nigra are lost [4–7]. In most cases, by the time PD is diagnosed, the disease may have already started 4 to 6 years beforehand [7]. So there is a need for developing a test for detecting PD *before* the onset of the motor symptoms (called Presymptomatic PD (PPD) [8], [9]). Such a test would help a clinician not only in detecting PPD, but also in monitoring progression of PD or in monitoring efficacy of early therapeutic interventions [10, 11]. Detecting PPD helps patients to start their treatment in the presymptomatic phase. It has been established that there are advantages of early pharmacological and therapeutic intervention in PD such as monoamine oxidase B inhibitors [12], catechol-o-methyl-transferase inhibitors [13], amantadine [14], amplitude training [15], reciprocal pattern training [16], and gait-balance training [17], including, a reduction in symptoms, particularly dyskinesia, and the delay of levodopa initiation [18]. Both the reduction of symptoms and the potential for slowing disease progression has a significant impact on improving patient quality of life [18].

Although there have been recent advances in identifying potential biomarkers including genetic [19, 20] and neuroimaging techniques [11] that help in detecting PD, these are still in early stages, and further work is needed in this direction. Further, it has also been recognized that a variety of nonmotor symptoms associated with PD may be observed years before the onset of motor symptoms and hence may be used as markers for detecting PPD [21]. Examples of these nonmotor symptoms are olfactory loss [22, 23], rapid eye movement sleep behavior disorders [24], bowel dysfunction and so on [21]. Recently, [7] proposed that a combination of tests (including smell test, transcranial sonography, and SPECT) could be helpful in detecting PPD. Further, [25] proposed a general methodology and an automatic system that can be used for detection of presymptomatic phase or diagnosis, and/or to monitor the treatment effectiveness for a variety of neurological disorders based on eye movement data to various stimuli. An important ingredient that is lacking in these methodologies is that these approaches are empirical and are not based primarily on an understanding of the mechanism causing these symptoms.

As widely noted in the literature [26–39], the sensorimotor system of a healthy individual may be seen as a stable control system, while the sensorimotor system of an individual with PD showing motor symptoms may be viewed as a control system with instabilities. The source and mechanism of these instabilities still remain topics of debate and investigation with delay-induced instability (with higher loop-delay in PD causing the instability, further explained in section 2.1) being one of the prominent theories [32, 36, 37, 39]. In general, the latencies in the sensorimotor loop are also known to be amplified and cause instabilities in other conditions as well like Multiple Sclerosis (MS) and stroke survivors [40]. In this paper, we propose a method for detecting this degradation in stability, which can then be applied to detecting PPD based on observing the progression of the tendency of the sensorimotor loop to develop instabilities *before* the instability manifests itself. Here, we do not concern ourselves with the source or mechanism of the instability and develop a method that could be utilised for any instability in general. This proposed method is further fleshed out through a numerical study. We employ a simple pendulum with PID controller as a conceptual representation of the sensorimotor loop to simulate motor control tasks in healthy and PPD individuals for this analysis. We also present an example task with physiological measurements that can be used with this method and can potentially be utilized in a clinical setting. We present representative human subject data collected through such a test, thereby demonstrating the feasibility to generate data for the proposed method.

The conceptual framework of the proposed method is described in detail in Section 2.2, the simulation example used for our numerical study in Section 2.3, and potential methods to classify whether a particular data is from healthy individuals or an individual with PPD in Section 2.3 and 2.3. The results of the numerical study are described in Section 3. An example task and associated data collected from human subjects is presented in Section 3.2. The longitudinal study needed to validate the method will be taken up as future work. Finally, we discuss further steps needed to validate the proposed method in Section 4 and close with the conclusion in Section 5.

## 2 Methods

In this section, we first describe a simple representation of the sensorimotor loop. Subsequently, we describe how we leverage the insight that in PD, the instability in the sensorimotor loop develops gradually over a period of time, to propose an approach for detecting PPD. Based on the proposed approach, we perform numerical simluations using a simplified representation of sensory-motor loop to generate synthetic data, and test the method using a numerical study.

### 2.1 Sensorimotor Loop Representation

One can represent the motor control (movement control) in humans as a simple feedback control system of the form shown in Fig. 1. In the schematic shown, the body dynamics is the natural dynamics in the absence of any neural control of the body part of interest (e.g. hand). The feedback path represents all sensory feedbacks including proprioceptive feedback, visual feedback, and tactile feedback. These sensory feedbacks are carried to the controller (brain) by afferent nerves. The controller represents the neurosystem’s logic that continuously compares the kinematic variables from sensory feedbacks (e.g. actual velocity) with the desired kinematic variables (e.g. desired velocity) to determine motor command. This motor command is then conveyed through the efferent nerves and results in muscle actions to get the desired response. We refer to this closed-loop feedback system consisting of motor actions and sensory feedbacks as the human sensorimotor system. As there are various time delays in the human sensorimotor system including delays due to nerve conduction times and information processing time, for simplicity, we lump all sensorimotor system delays (delays in various portions of the sensorimotor system) into one transport delay in the closed-loop feedback system. Finally, the saturation in the loop approximates the physiological limit of the transmission of neural control actions [41].

**Figure 1.**
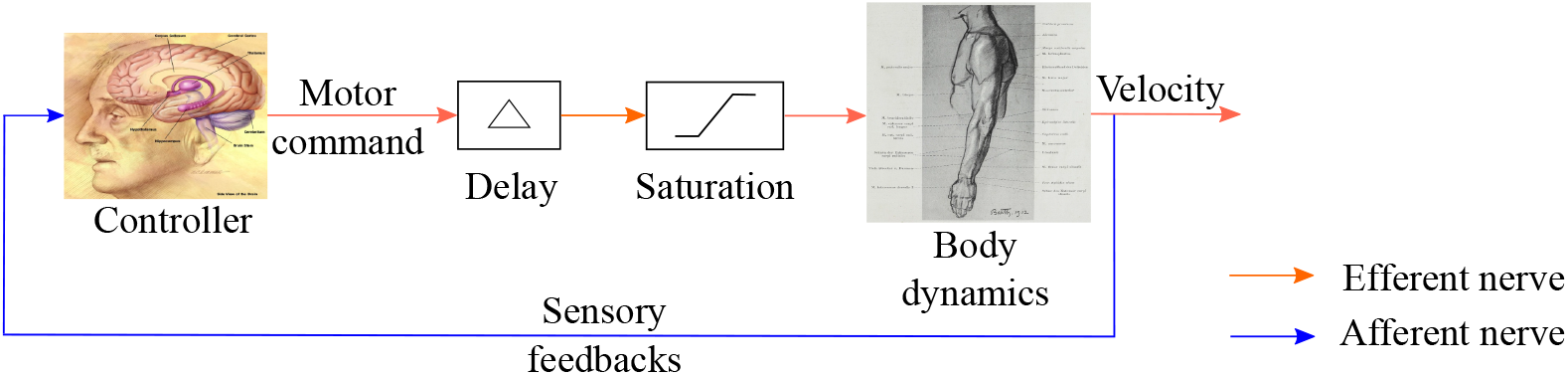
The closed-loop feedback system representing the human sensorimotor system

A number of studies [26–28,32,35,37–39,42] have observed that increased sensorimotor loop delay in PD tends to destablize the sensorimotor system. This is supported by the observation that response time in patients with PD is larger compared to healthy individuals [43–46]. According to these observations, the sensorimotor system of a healthy individual is a control system with a smaller delay and the sensorimotor system of an individual with PD is a control system with a larger delay. Thus, this larger delay, that is, a delay beyond a certain threshold, leads to an instability in the human sensorimotor system. This observation also explains some of the clinical features of parkinsonian rest tremor [37, 39] and provides a possible explanation of how high-frequency deep brain stimulation suppresses low-frequency rest tremors [47]. There are some works that also consider other possibilities for the source of instabilities including an increased gain in the sensorimotor loop [32]. However, as stated earlier, the subsequent development will not rely on this assumption but simply focus on the degradation of stability and will be applicable regardless of the source of instability.

With this background, in Section 2.2, we provide the main concept of the proposed methodology.

### 2.2 Proposed Approach for Detecting Presymptomatic Parkinson’s Disease (PPD)

Based on the insight of degrading stability in the sensorimotor loop, the proposed approach hinges on the detection of the transition of the human sensorimotor system from a stable system (healthy individual) to an unstable system (individual with PD) on the basis of response recorded from a simple movement control task in the clinic and repeated several times over a period of time. Note that we expect this transition of the human sensorimotor system from a stable to an unstable system to be gradual as PD is a slowly progressing disease.

In control system theory, the *poles* of a linearized system are a set of complex numbers that represent the nature of (and more specifically the exponents associated with) the transient behavior and thus stability of that system. The poles of the dynamical system are also roots of differential equation that governs the system’s dynamics. A system with poles in left-half of the complex plane is stable, that is, transient response decays with time, whereas a system with at least one pole in right-half of the complex plane is unstable, that is, transient response grows with time. We therefore propose to use the poles as a measure of stability of the representative human sensorimotor system involved in a specified clinical task.

The proposed approach can be summarised as follows, a subject performs a series of a specific clinical task on several occasions repeated over a period of time (not necessarily at equal intervals) and poles are estimated from these data using an algorithm (explained in Section 2.3). Finally, a classification algorithm is applied to the estimated poles to classify whether a particular individual is healthy or has PPD.

A degradation of stability implies that at least one of the poles is gradually moving towards the right-half of the complex plane. Hence, to observe the gradual movement of pole(s), it is required to keep track of pole(s) over a period of time rather than observing pole(s) at a particular time instant, and consequently subjects have to repeat the same clinical movement control tests on several occasions over a period of time (e.g. as part of a routine yearly checkup). Significant movement towards right half of complex plane of at least one of the estimated poles from these clinical tasks indicates PPD.

We now consider a simple simulation example to further detail out the proposed method.

### 2.3 Simulation Example

For our numerical exploration, we use a simulation example as shown in Fig. 1, we use a simple pendulum to represent the dynamics of the task and a Proportional-Integral-Derivative (PID) as a controller, along with a delay and saturation in a unity feedback configuration. The simple pendulum has length *L*, mass *M*, and the damping coefficient *C*. For our simulation purpose, we use *L* = 0.65 *m, M* = 3.5 *Kg, C* = 3.375 *Kgm/s*, and a PID controller with the Proportional gain *k*_*p*_ = 15, Integral gain *k*_*i*_ = 4 *Hz* and Derivative gain *k*_*d*_ = 0.5 *s*. All simulations were carried out using MATLAB & Simulink of MathWorks (Natick, Massachusetts).

This simulation example is similar to a movement control task in a clinic and has the same structure as shown in Fig. 1 and similar to other examples in the literature [26, 32, 37, 39, 47]. To simulate degradation of stability, we first consider a small and constant delay as a representation of the human sensorimotor system of a healthy individual, and a gradually increasing delay as a representation of the human sensorimotor system for an individual with PPD. Note that this gradual increase in the delay is still below the delay threshold, that is, the delay has increased but not to the extent that it destabilizes the human sensorimotor system leading to visible motor control symptoms. Similar to delay analysis, to demonstrate that the method works equally well for other mechanisms of instability, we subsequently also consider a constant gain as a representation of the human sensorimotor system of a healthy individual and a gradual increment in gain as a representation of the human sensorimotor system for an individual with PPD. Analogous to delay, the gain is increased but not to the extent that it destabilizes the human sensorimotor system.

### 2.4 Procedure for generating synthetic data set

As explained in Section 2.2, a subject needs to repeat the same clinical task on several occasions over a period of time. To mimic within-subject variabilities in the sensorimotor delay values (or gain values) over a period of time due to various physiological factors, we consider an additional stochasticity around the constant value of the delay (or gain) representing healthy individuals and a similar stochasticity around a gradual increasing delay (or gain) value representing PPD individuals. This stochasticity is assumed to follow a Gaussian distribution. Further, to represent noise in the human sensorimotor system, we introduce system and measurement noise with zero mean and Gaussian distribution.

Next, we use these sensorimotor delay values (or gain values) in the simulation example to generate simulated responses similar to data from a clinical task and generate several instances of the same task repeated over a period of time.

As an example, Fig. 2 shows a simulated data of a single clinical task. Following this procedure, we generate a synthetic data set simulating 500 instances of sensorimotor loop system for healthy individuals and 500 instances of sensorimotor loop system with a gradual increment in gain and delay representing PPD. This synthetic data represents 1000 individuals for training and validation needed for the machine learning algorithm. We used 600 individuals (synthetic data set) for training and 400 individuals (synthetic data set) to validate the machine learning algorithm. The numerical values of delays and gains for generated synthetic data set are given in Appendix 6.1.

**Figure 2.**
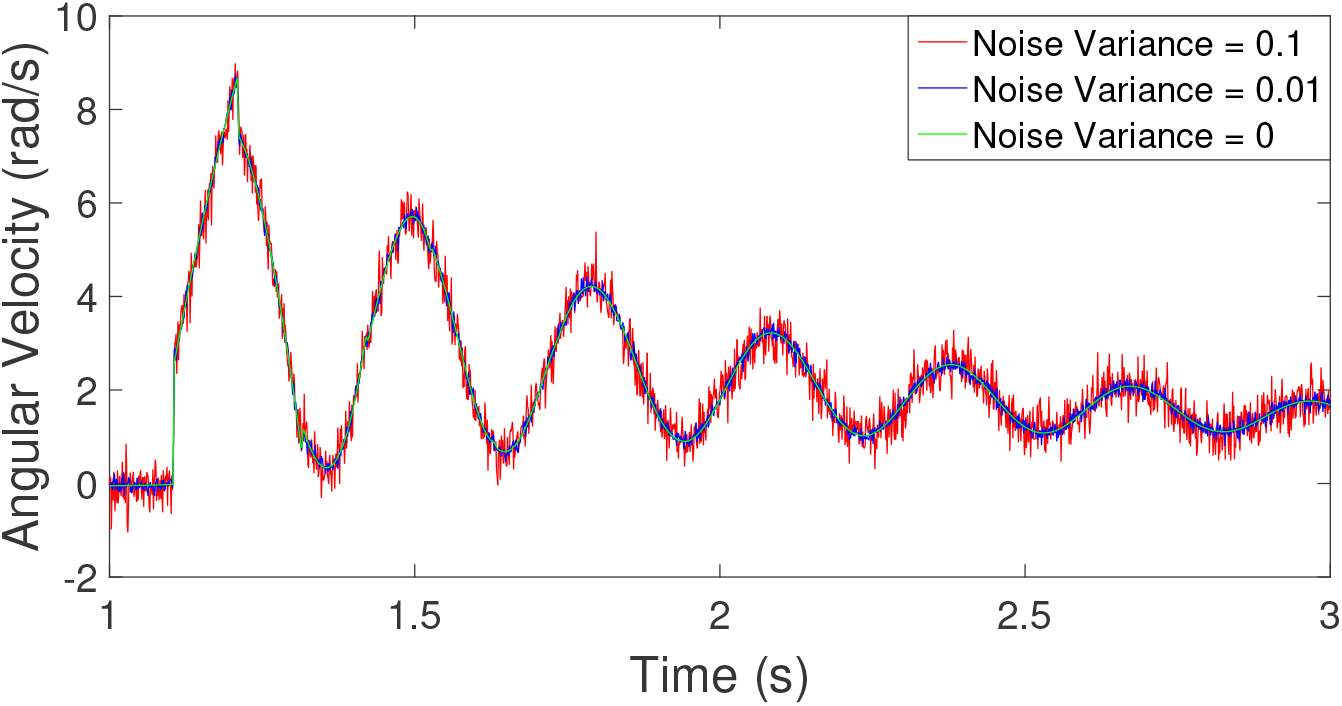
Output of simulation example representing data from a single clinical movement control test with different variances of noise. Here the sensorimotor delay value is 0.105 *s* with the addition of system and measurement noise of variance 0, 0.01 and 0.1, respectively

### 2.5 Estimation of poles from data

To estimate the poles from data, we propose to use a Matrix Pencil Method (MPM) [48]. MPM approximates time series data by *M* complex exponentials. These estimated complex exponents represent the poles of the system of interest (in our case, the human sensorimotor system). The observed signal is represented as

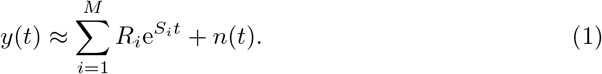

Here, *y*(*t*) is simulated response, *S*_*i*_ are the estimates of poles, *R*_*i*_ are coefficients and *n*(*k*) is the noise in the simulated data. A brief overview of an algorithm for estimating poles from data is given in Appendix 6.2.

Fig. 3 shows real part of poles for healthy and PPD individuals over a period of time, estimated from a few individuals in the synthetic data generated as described earlier. Further, it is to be noted that the synthetic data set consists of both individuals maintaining a regular frequency of visits and individuals maintaining an irregular frequency of visits, that is, the clinical tasks are not necessarily repeated at regular intervals but could be at irregular intervals.

**Figure 3.**
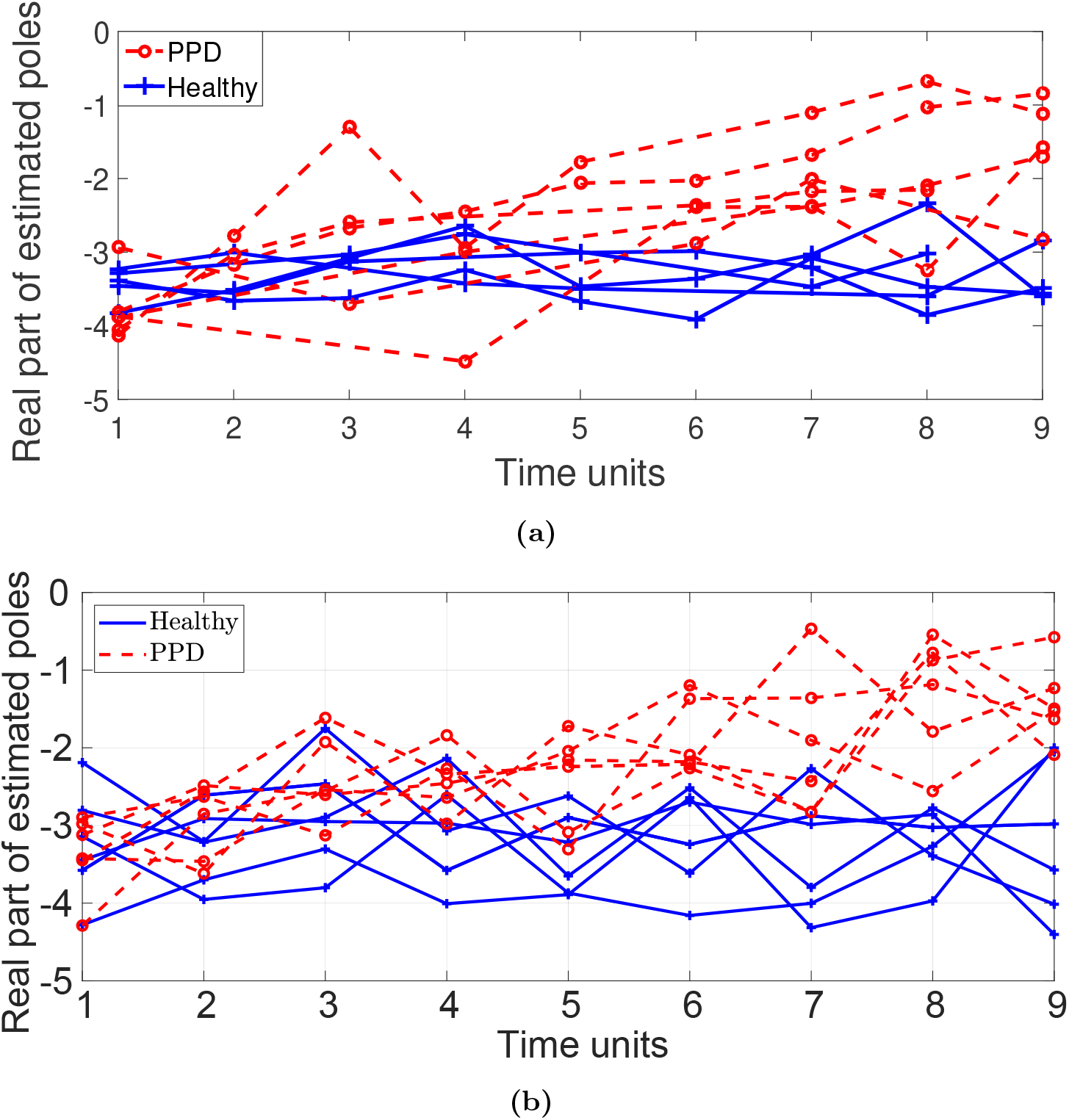
The real part of estimated poles from the synthetic data set representing sensorimotor loop system for ten individuals over nine-time units (each unit may represent several months) for a) gradual increment in delay, and b) gradual increment in gain. Out of 10 individual data, 5 represent healthy individuals (solid blue), and 5 represent PPD (dashed red). These data also show the regular or irregular nature of clinical movement control tests over time units. Simulations with gradual increment in gain are at regular intervals, and simulations with gradual increment in delay are with regular and irregular intervals of clinical movement control tests. It is clear that due to the stochasticity, the trends are not clearly distinguishable (especially if there are only a few data points), and hence a robust classification algorithm is needed

### 2.6 Machine learning algorithm for classification

Once we estimate poles from the measured data, the next step is to apply a classification algorithm for classifying whether a particular data (for an example, one of the traces in Fig. 3) is from data-set representing healthy individual or data-set representing an individual with PPD. In healthy individual data-set, despite variabilities due to numerous factors, we do not expect the estimates of poles to have a significant trend of moving towards right-half of the complex plane. On the other hand, for the data-set representing individuals with PPD, we expect at least one of the estimates of poles to have a significant trend of moving towards right-half of the complex plane. Now, one way to identify this significant trend compared to healthy individuals is to use simple statistical tests or a classifier based on simple thresholds. Exploring this approach with the synthetic data, we find that a reliable classification using a threshold alone is not a robust approach. This is partly due to the significant overlap between poles of the human sensorimotor system for healthy and PPD groups in simulated data (representing inter subject variability), and perhaps partly due to the stochastic variability as apparent in Fig. 3. Therefore, we explore machine learning algorithms for a more robust classification. Typically, the input to the machine learning algorithm is a set of features derived from the data. For instance, to classify whether a particular individual has a benign or malignant tumor, the tumor size and age at diagnosis are possible features to consider [49]. Using these features, a machine learning algorithm trains a model that best describes input (features)-output (class) relationship for those data and uses this model to classify the new data. In the next subsections, we briefly describe two machine learning algorithms.

1. Support Vector Machine: Support Vector Machine (SVM) [50] is a supervised machine learning algorithm that given the training data with their features and class labels identified a priori, determines the maximum margin hyperplane in feature space. Maximum margin hyperplane is a plane in feature space from which distance to the nearest data points of both classes is maximized. Once maximum margin hyperplane is determined, depending upon the position of the new data set in feature space, that is, whether a new data is lying below the hyperplane or above the hyperplane, each new data set is classified as either Class 1 or Class 2. In the case of outliers in data or not linearly separable data, a variant of SVM is used that finds a non-linear boundary to separate both classes of data. For further details, refer to [50]. **Feature Selection:** Feature selection is an important aspect for the success of machine learning algorithms. A good choice of features helps improve classification performance, lower computational complexity, build better generalizable models and decrease required storage [51]. The aim of feature selection is to extract features from data that represent characteristics of each of the class or groups. Since we are interested in tracking the stability of the human sensorimotor system and only the real part of estimated poles in the complex plane governs the stability, we only use the real part of estimated poles for the rest of our analysis. We explore two sets of choices (C1 and C2) for feature selection. These choices of features are then used as input to the SVM to classify whether a particular individual is healthy or has PPD. **C1:** Since we are interested in the trend of the real part of the estimated poles over a period of time, percentage change in the real parts of poles with respect to the baseline (the estimated poles from the simulated response of the first movement control test), and percentage change in the successive difference in the real parts of poles between simulated responses over a period of time are the features worth considering. But, in reality, it is very likely that the clinical task is not performed at fixed intervals of time for all subjects. Hence, we consider the rate of percentage change (either per month or per year) for both of the above mentioned quantity as features. Based on preliminary simulation results, we find that only three features are sufficient for robust classification, namely; minimum of percentage change rate in the real part of the poles, maximum of percentage change rate in the successive difference between tests and real part from the first clinical movement control test. The calculation of percentage change rate in the real part of the poles and percentage change rate in the successive difference between tests are given in the Table 1. Here, *N* is the total number of clinical movement control test. that are conducted possibly at irregular time intervals. *T*_*k*_ is the time duration between two trials, where *k* = 1, 2, …, *N* − 1. *x*_1_ to *x*_*N*_ are real part of estimated poles with subscripts indicating the test number. **C2:** For a second choice of features, since we are interested in determining whether the real part of estimated poles has an increasing trend, we use hypothesis testing as a tool to detect a statistically significant increasing trend in presence of noisy data. In this approach, we use the statistical value of slope and constant of the real part of estimated poles as features for the classification algorithm. Further, we also take the real part of estimated poles of the simulation response of the first movement control test as the third feature as an indicator of the baseline for each individual.
2. Support Vector Machine for Longitudinal Analysis (LSVM) A recently developed method called Longitudinal Support Vector Machine (LSVM) [52] is a method specifically developed for longitudinal data. Here, each data point takes the form of a single time-series. LSVM is shown to have higher accuracy compared to SVM, linear discriminant analysis (LDA), functional linear discriminant analysis (FLDA). Note that once we have real parts of estimated poles over a period of time, this method is formulated such that an additional feature selection is not required.

## 3 Results

In this section, we first test the robustness of MPM and discuss the classification results of two machine learning algorithms on the synthetic data set. First, to test the robustness of the MPM in the presence of noise, we introduce system and measurement noise with different variances. Fig. 4 shows an estimate of poles for various values of delay and gains and the effect of noise with different variances on estimation of poles, indicating that the estimates from MPM are robust to noise and disturbances. For all the data generation, we assume that system and measurement noises have the same variances.

**Table 1.**
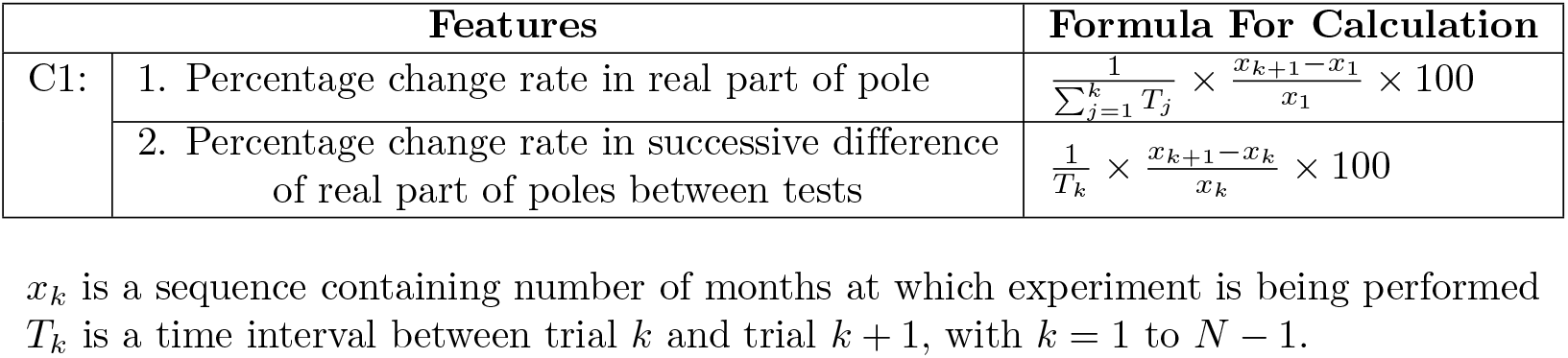
Calculation of features for *C*1 from the estimated poles

**Figure 4.**
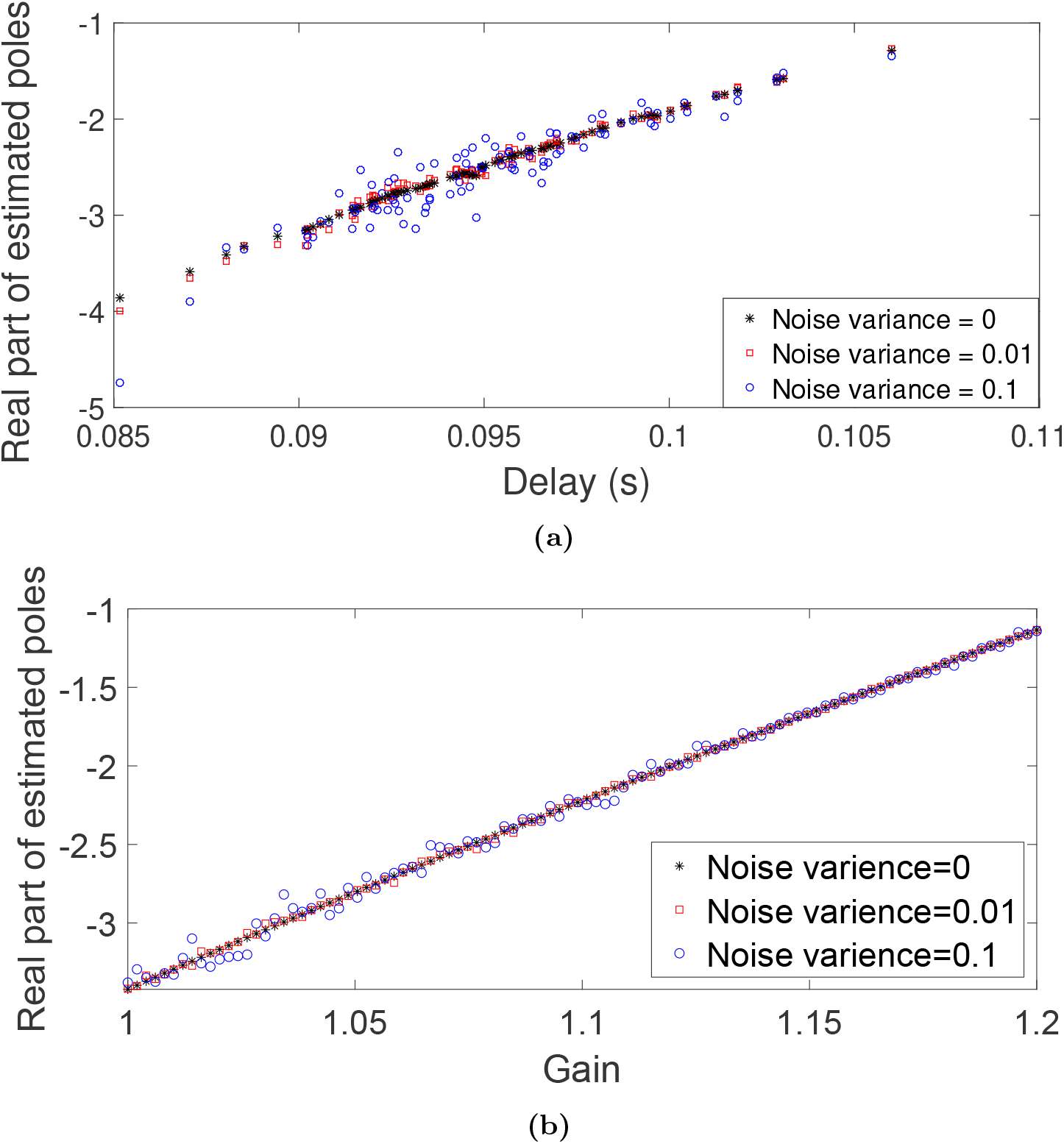
Effect of noise variance on the estimation of poles using MPM, (a)change in poles due to change in delay (b) change in poles due to change in gain

### 3.1 Classification Results

Fig. 5 shows a box-plot of the real part of estimated pole for the synthetic data set containing healthy and PPD group, over a period of time. The box-plot shows the median, range and the quartiles of the population distributions. It is clear from the box-plot that real part of estimated poles remains in the same range for healthy individuals and increases for PPD over a significant period of time. Next, we apply machine learning algorithms to these data.

**Figure 5.**
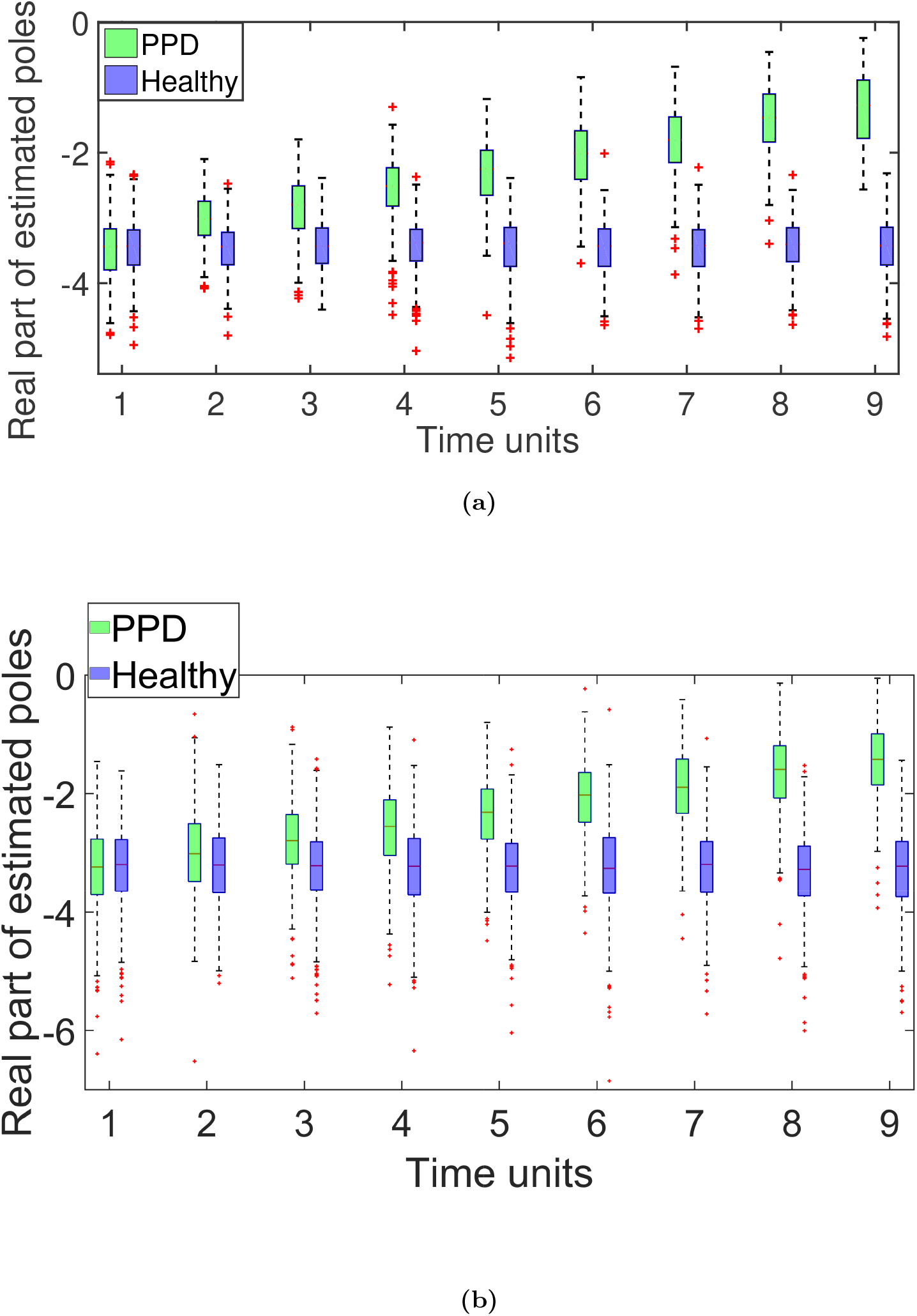
Box-plot of a synthetic data set (total 1000 individuals, 500 individuals in each group) of the real part of estimated poles for 9 time units (each unit may represent several months) for healthy and PPD for a) gradual increment in delay, and b) gradual increment in gain. It is observed that the real part of estimated poles remains in the same range for healthy individuals and increases for PPD over a significant period of time.

To show the efficacy of the proposed method, we compare our results of SVM (incremental delay) with proposed features (C1 and C2) with the results of the LSVM (incremental delay). The classification results for simulated (incremental delay) data set with a noise variance of 0.01 are shown in Table 2 and a noise variance of 0.1 are shown in Table 3. It is seen from the Table 2 and 3 that the results of SVM with features C2 are comparable with the results of LSVM. It is clear from Table 3 that the above results are almost unaffected by a noise variance of 0.1. Further, on repeating the above procedure with ten independently generated synthetic data sets, we find that the above results are reproducible. Note that all of the methods have a high sensitivity of above 96% and specificity above 98%. In addition, they perform very well with the validation synthetic data set with sensitivity above 95% and specificity above 98%. All three approaches seem to perform well with this synthetic data set.

**Table 2.** Comparison of classification result of synthetic (incremental delay) data set with disturbance and measurement noise of zero mean and 0.01 variance with the choice of features for C1 being minimum of percentage change rate in real part of characteristic value and maximum of percentage change rate in successive difference and the choice of features for C2 being statistical value of slope and constant of straight line fit to data and real part of characteristic value of first trail as features.

| Methods | Training synthetic data set |  |  |  |  |
| --- | --- | --- | --- | --- | --- |
|  | Sensitivity (%) | Specificity (%) | False Positive (%) | False Negative (%) | Error in Classification (%) |
| **LVSM** | 96.73 | 100 | 0 | 3.27 | 1.67 |
| **SVM (C1)** | 96.40 | 98.97 | 1.03 | 3.60 | 2.34 |
| **SVM (C2)** | 98.36 | 99.66 | 0.34 | 1.64 | 1 |
| Validation synthetic data set |  |  |  |  |  |
| **LVSM** | 97.42 | 100 | 0 | 2.58 | 1.25 |
| **SVM (C1)** | 95.36 | 99.51 | 0.49 | 4.64 | 2.5 |
| **SVM (C2)** | 98.96 | 99.02 | 0.98 | 1.04 | 1 |

**Table 3.** Comparison of classification result of simulated (incremental delay) data set, same as Table 2 except for the variance being 0.1.

| Methods | Training synthetic data set |  |  |  |  |
| --- | --- | --- | --- | --- | --- |
|  | Sensitivity (%) | Specificity (%) | False Positive (%) | False Negative (%) | Error in Classification (%) |
| **LVSM** | 96.65 | 100 | 0 | 3.35 | 1.67 |
| **SVM (C1)** | 96.66 | 98.33 | 1.67 | 3.34 | 2.5 |
| **SVM (C2)** | 97.65 | 99.35 | 0.65 | 2.35 | 1.5 |
| Validation synthetic data set |  |  |  |  |  |
| **LVSM** | 96.51 | 100 | 0 | 3.49 | 1.75 |
| **SVM (C1)** | 96.02 | 97.99 | 2.01 | 3.98 | 3 |
| **SVM (C2)** | 98.01 | 97.49 | 2.51 | 1.99 | 2.25 |

Next, to demonstrate that the same method is effective when the source of instability is different from an increased delay, we consider the synthetic data set with a gradual increasing gain (as mentioned in Appendix 6.1). With the C1 choice of features and SVM method, we observe that the Sensitivity is 96% and specificity is 94.33% with the training synthetic data set (incremental gain), while for the validation synthetic data set (incremental gain), the sensitivity is 98% with a specificity of 93%.

### 3.2 Example Task and Physiological Measurement

One of the major requirements in designing a clinical movement control test for detecting PPD is that the test should be simple and easy to execute. There are various possible clinical movement control tests such as; spiral tracing task in which the subject needs to trace a spiral drawn on white paper [53], eye tracking task in which the subject needs to follow a circle displayed on screen [54, 55], compensatory tracking task in which the subject needs to maintain the actual position of the arm with zero reference (or initial) position of arm displayed on the screen in presence of outside influence by means of compensating the error in arm position [56]. In this work, we consider pupil dilation/constriction data for applying the proposed method. A significant latency is reported during pupillary light reflex in patients with Parkinson’s disease [57, 58], and this latency increases with the progression of the disease [59]. This latency degrades the stability of the sensory-motor loop representing pupillary-light-reflex.Therefore the proposed method is applicable in this situation. It is worth noting that the proposed method is equally applicable to any of the other motor control tasks mentioned above. We developed a pupilometer in the lab as shown in Figure 6a that uses a camera, infrared light sources, LEDs to stimulate the pupil (with light flashes). With this pupilometer with its onboard microcontroller, it is possible to produce different stimulation patterns of white LEDs switching on and off, and the resulting pupil response is recorded as a video. Further, using relatively straightforward image processing techniques, the pupil diameter as a function of time is estimated from the recorded video. This is a simple portable device and a comfortable task for patients and can be conducted in any clinical environment without any special arrangements. A sample screenshot of the video recorded by the device after applying image processing to detect the pupil diameter is shown in Figure 6b.

**Figure 6.**
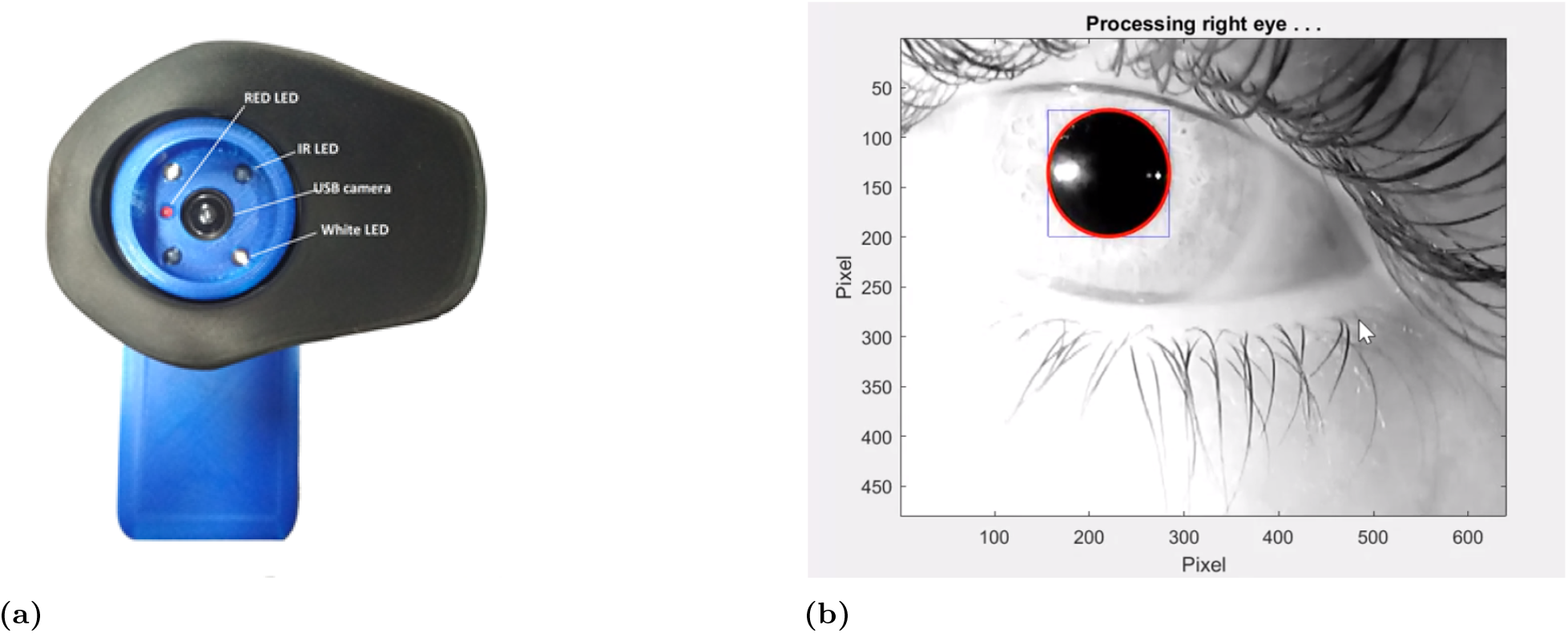
(a) The pupilometer device developed in the lab that is used to collect experimental data of pupil constriction/dilation dynamics that can potentially be used as a clinical test along with the proposed method, (b) Screenshot of the recorded video after image processing to identify and estimate the pupil diameter.

This device is used to collect the data from age-matched healthy control subjects and patients with PD. The Institute’s ethics committee approved the experimental protocols, and the participants gave their written informed consent before participating in the experiment. For all our tests, subjects were asked to focus on the red LED (center of a screen) during the test, with the LED light source being on for 10 second and off for 10 seconds. We repeat the same procedure 5 times in a single trial and performed three sets of trials. Pupil diameter data from each of these trials is essentially a step response and matrix pencil methods is used to estimate poles representing the motor control dynamics.

The output of the pupil detection algorithm for a few representative PD patients (6) and age-matched healthy individuals (6) are shown in Figure 7. A total of 63 trials were conducted and MPM was used to detect poles for all these trials, the pole data is not reported here as the absolute values are irrelevant and only long-term trends are relevant. The data shows feasibility of setting up a simple clinical task and ability to generate good quality data that the MPM is able to process to generate estimates of poles. The machine-learning-based classification algorithm is however not applied here as that will require long term data from a longitudinal study. The proposed method is designed to detect long-term trends in a single patient’s data and therefore comparison across patients is not of value and hence the quantitative results in Figure 6b are not discussed in detail. A longitudinal study to validate the method is planned as a future work.

**Figure 7.**
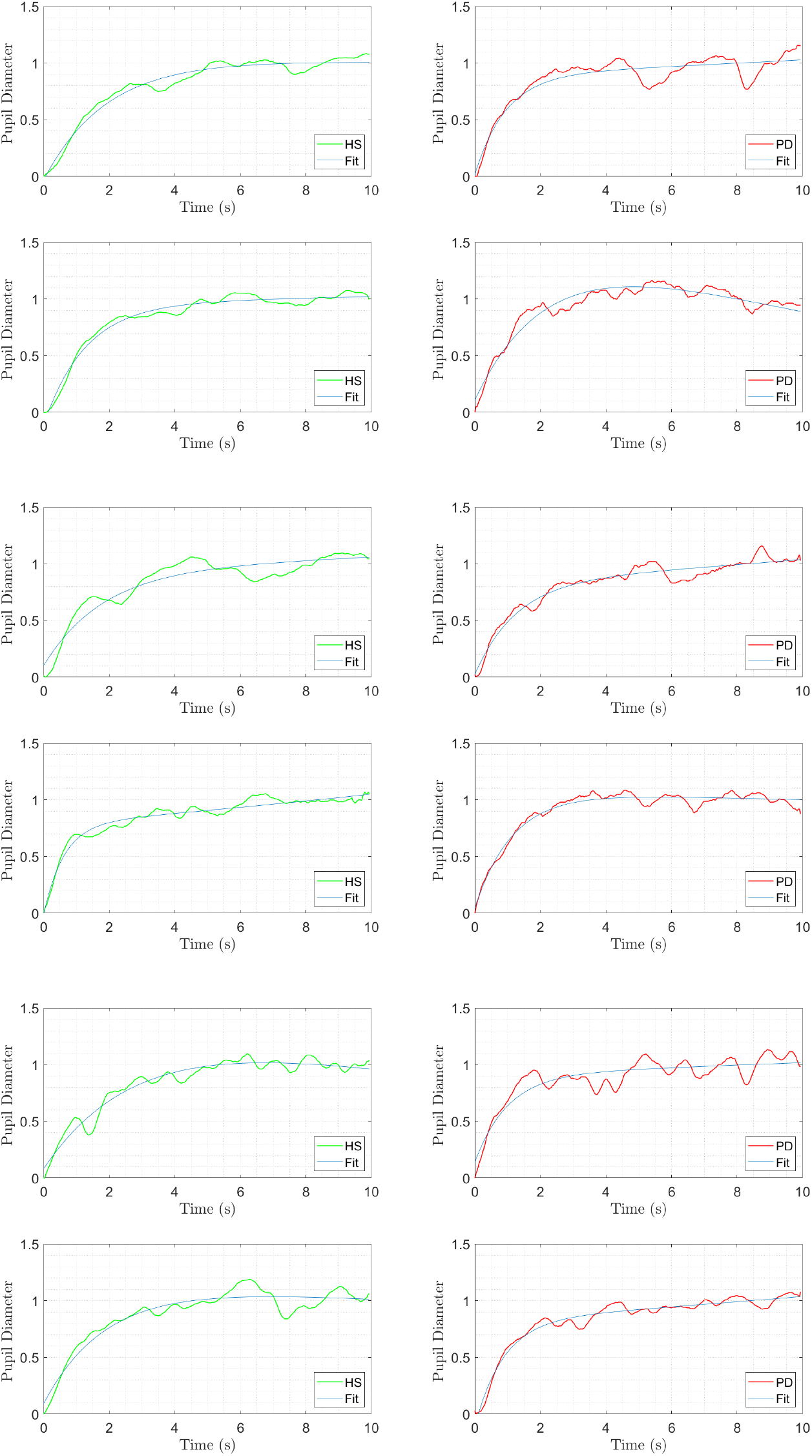
Plots of normalized pupil diameter as a function of time for both age-matched healthy subjects (green) and PD patients (red). The plots shows one subtrail each for 12 subjects with pupil diameter increasing when LED in the pupilometer device is switched off. The data is plotted after a moving-average filter is applied to smooth out the noise. The MPM method is applied to this data to estimate the poles. The machine-learning-based detection algorithm can however not be applied at the moment as that algorithm is meant to detect long-term trends with a longitudinal data set.

## 4 Discussion

In continuation of the proposed approach and simulation results presented in this paper, a longitudinal study with real patient data is needed to further analyse the effectiveness and robustness of the methods. The proposed methodology appears to show potential, and future tests will help develop this idea into a viable early diagnosis device. The proposed methodology has potential applications in other situations including motor performance in multiple sclerosis [60], stability of balance control [61, 62], characterizing slow eye movements [63], wearable health monitoring [64] and cerebral autoregulation [65, 66].

The sequence of steps required to implement the proposed approach in a clinical setting are described in Fig. 8. In summary, a subject performs a series of clinical movement control tests on several occasions over a period of time (not necessarily at equal intervals) and each data is recorded. After a few clinical movement control tests spanning a significant amount of time (say several months or years), using an algorithm (explained in Section 2.3), poles are estimated from these data for a particular individual. Finally, a classification algorithm is applied to the estimated poles. The output of the classification algorithm classifies whether a particular individual is healthy or has PPD. This classification will directly aid clinicians identify individuals who are at risk of developing PD in near future and take necessary precautions or apply certain interventions as appropriate.

**Figure 8.**
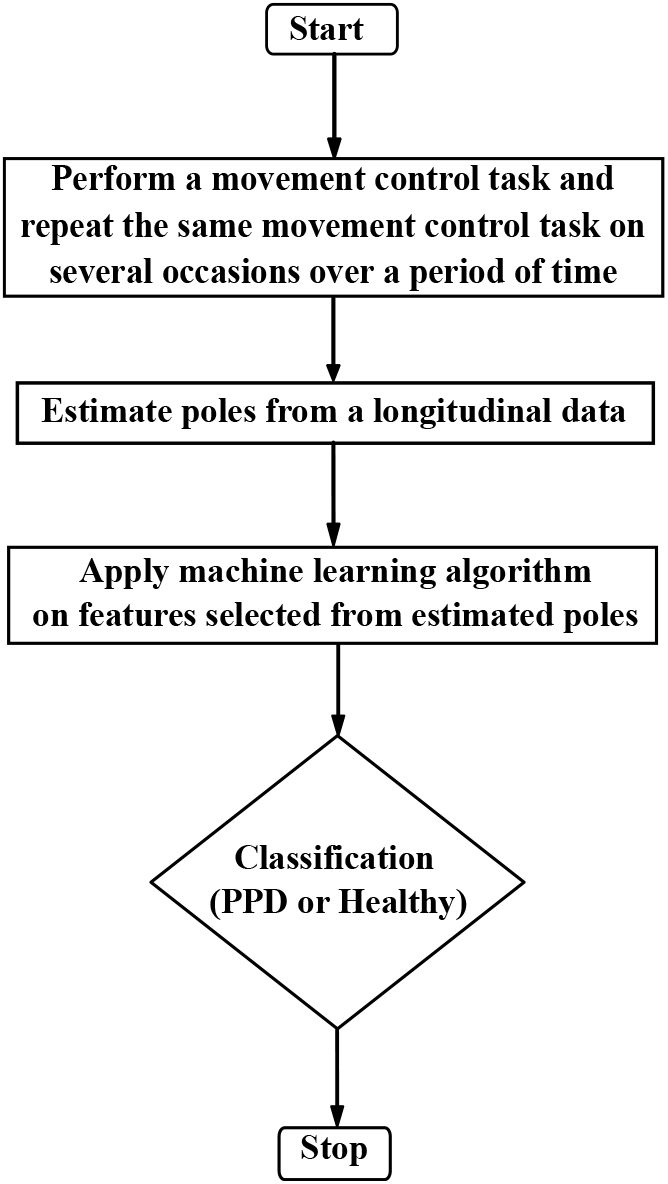
Flowchart of a proposed methodology for detecting PPD in a clinical setting

One possible clinical task with a simple pupilometer outlined in Section 3.2 can be utilized for generating data. Further, one can also choose other clinical tasks as discuss earlier with a step input (step change) as this is easy to implement while at the same time yielding a response that can directly be used to estimate the poles of the system. An example of a step input is a step change in circular target position, that is, considering a circular target initially at some arbitrary position and instantly the target moves to another position and stays there.

## 5 Conclusions

In this paper, we proposed a novel methodology for detecting degradation of stability in the sensorimotor loop that is applicable to detection of PPD even before the appearance of clinical symptoms in PD. This proposed method is based on the key insight that the gradual development of motor symptoms in PD can be seen as a gradual degradation in stability of the senosriomotor loop, and the fact that the location of poles of a closed-loop system in the complex plane characterizes the stability of the system. Therefore, the key idea is to detect the gradual progression of the stability of the human sensorimotor system (before the system actually becomes unstable) from experimental data collected by performing a simple clinical movement control test on several occasions over a period of time. The proposed method is evaluated on a synthetic data set and is seen to show promise and potential for use for detecting PPD through an early diagnostic device. An example task with physiological measurement that can potentially be used as a clinical movement control test along with representative data was also presented demonstrating the feasibility of performing a longitudinal study to validate and test the robustness of the proposed method.

## 6 Supporting information

**S1 Appendix**.

### 6.1 Numerical Simulation of Data Set using Simulation Example

To mimic the variabilities in the delay values due to various physiological conditions over a period of time, we choose the constant value of delay to be 0.088 s and consider small stochasticity around the constant value of the delay for healthy individuals and small stochasticity around the gradual increase in delay values (from 0.088 s to 0.11 s) for PPD individuals. These stochastic variabilities follow Gaussian distribution and are generated such that signal to noise ratio is 30 dB. To simulate instability due to gain, we choose a gradual increment in gain with a 0 to 20% increase in controller gains (*k*_*p*_ from 15 to 18, *k*_*i*_ from 4 to 4.8 *Hz* and *k*_*d*_ from 0.5 to 0.6 *s*) with the small stochasticity around constant 0.088s delay. Further, we introduce a system and measurement noise (with zero mean and 0.01 variance) in the simulation example. Next, we run the simulation with a step input for each value of the delay and gains. From the step response data, we estimate poles using MPM. We follow the same procedure for all values of delay and gains to get a real part of estimated poles over a period of time for each of the individuals. Fig. 3 shows an example of a simulated data set of the real part of estimated poles for healthy and PPD individuals over time.

### 6.2 Matrix Pencil Method (MPM) [48]

The MPM approximates a time series data by a sum of complex exponentials as

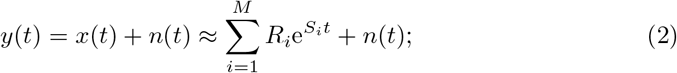

where,

*y*(*k*)= observed time response data

*x*(*k*)= original signal

*n*(*k*)= noise in the system and measurement

*R*_*i*_ = coefficient and

*S*_*i*_ = −*σ*_*i*_ + *jω*_*i*_ = poles.

Due to noise in the data, we used a total-least-squares MPM. In this, we form a data matrix as,

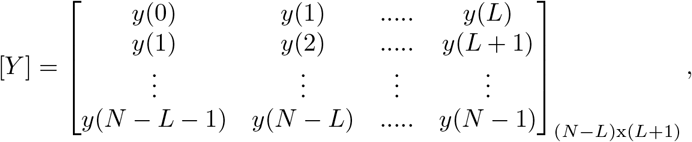

where *L* is noise filtering parameter and for a efficient filtering, it should be chosen between *N/*3 to *N/*2. Next, we find a singular value decomposition (SVD) of the matrix [Y]

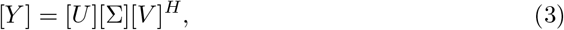

where, the superscript *H* denotes the conjugate transpose. To select *M*, we see the ratio of the various singular values to the largest one. Generally, we consider the singular values *σ*_*c*_ such that 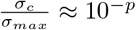, where *p* is the number of significant decimal digits in the data. So *M* is the value at which above ratio is greater than equal to 10^*−p*^. Now, we construct a matrix, [*V* ′] which is called a “filtered” version of [*V*] and it contains only M dominant right-singular values of [*V*]. Therefore,

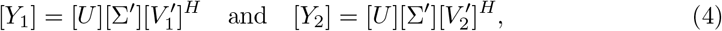

where, 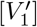 is obtained by deleting the last row of [*V* ′] and 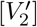 is obtained by deleting the first row of [*V* ′]. [Σ′] is obtained from *M* column of [Σ] corresponding to M dominant singular values. Finally, the poles (*z*_*i*_) can be obtained by solving the generalized eigenvalue problem of the following matrix

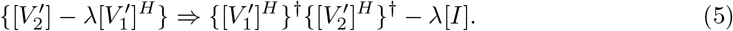

Once, *z*_*i*_ are known, the residue *R*_*i*_ are solved from the following least-square problem

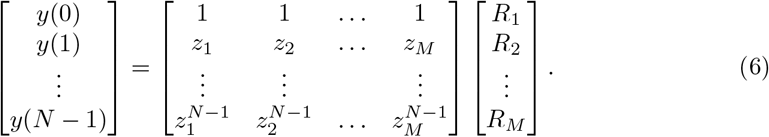

## 7 Acknowledgments

The authors are grateful to the people of BKP-Parkinsons’s Disease and Movement Disorders Society (BKP-PDMDS), senior faculty members of IIT Gandhinagar, Dr Dhvani Parikh and Dr Jitender Singh, for helping us during data collection. We also acknowledge Prof. Seth Blumberg (University of California, San Francisco) for helpful discussions.

## Notes

### Competing Interest Statement

The authors have declared no competing interest.

## References

1. Sharma S, Moon CS, Khogali A, Haidous A, Chabenne A, Ojo C, et al. Biomarkers in Parkinson’s disease (recent update). Neurochemistry international. 2013;63(3):201–229.

2. Jankovic J. Parkinson’s disease: clinical features and diagnosis. Journal of Neurology, Neurosurgery & Psychiatry. 2008;79(4):368–376.

3. Rao G, Fisch L, Srinivasan S, D’Amico F, Okada T, Eaton C, et al. Does this patient have Parkinson disease? Jama. 2003;289(3):347–353.

4. Mahlknecht P, Seppi K, Poewe W. The concept of prodromal Parkinson’s disease. Journal of Parkinson’s disease. 2015;5(4):681–697.

5. Fearnley JM, Lees AJ. Ageing and Parkinson’s disease: substantia nigra regional selectivity. Brain. 1991;114(5):2283–2301.

6. Greffard S, Verny M, Bonnet AM, Beinis JY, Gallinari C, Meaume S, et al. Motor score of the Unified Parkinson Disease Rating Scale as a good predictor of Lewy body–associated neuronal loss in the substantia nigra. Archives of neurology. 2006;63(4):584–588.

7. Sommer U, Hummel T, Cormann K, Mueller A, Frasnelli J, Kropp J, et al. Detection of presymptomatic Parkinson’s disease: combining smell tests, transcranial sonography, and SPECT. Movement disorders. 2004;19(10):1196–1202.

8. Postuma R, Montplaisir J. Predicting Parkinson’s disease–why, when, and how? Parkinsonism & related disorders. 2009;15:S105–S109.

9. Wu Y, Le W, Jankovic J. Preclinical biomarkers of Parkinson disease. Archives of neurology. 2011;68(1):22–30.

10. Miller DB, O’Callaghan JP. Biomarkers of Parkinson’s disease: present and future. Metabolism. 2015;64(3):S40–S46.

11. Sherer TB. Biomarkers for Parkinson’s disease. Science translational medicine. 2011;3(79):79ps14–79ps14.

12. Dezsi L, Vecsei L. Monoamine oxidase B inhibitors in Parkinson’s disease. CNS & Neurological Disorders-Drug Targets (Formerly Current Drug Targets-CNS & Neurological Disorders). 2017;16(4):425–439.

13. Rivest J, Barclay CL, Suchowersky O. COMT inhibitors in Parkinson’s disease. Canadian journal of neurological sciences. 1999;26(S2):S34–S38.

14. Rascol O, Negre-Pages L, Damier P, Delval A, Derkinderen P, Destée A, et al. Utilization patterns of amantadine in Parkinson’s disease patients enrolled in the French COPARK study. Drugs & Aging. 2020;37(3):215–223.

15. Farley BG, Koshland GF. Training BIG to move faster: the application of the speed–amplitude relation as a rehabilitation strategy for people with Parkinson’s disease. Experimental brain research. 2005;167(3):462–467.

16. Irwin-Carruthers S. An approach to physiotherapy for Parkinson’s disease. South African Journal of Physiotherapy. 1971;25(1):5.

17. Radder DL, Lígia Silva de Lima A, Domingos J, Keus SH, van Nimwegen M, Bloem BR, et al. Physiotherapy in Parkinson’s disease: a meta-analysis of present treatment modalities. Neurorehabilitation and neural repair. 2020;34(10):871–880.

18. Murman DL. Early treatment of Parkinson’s disease: opportunities for managed care. The American journal of managed care. 2012;18(7 Suppl):S183–8.

19. McInerney-Leo A, Hadley DW, Gwinn-Hardy K, Hardy J. Genetic testing in Parkinson’s disease. Movement disorders. 2005;20(1):1–10.

20. Tan EK, Jankovic J. Genetic testing in Parkinson disease: promises and pitfalls. Archives of neurology. 2006;63(9):1232–1237.

21. Haas BR, Stewart TH, Zhang J. Premotor biomarkers for Parkinson’s disease-a promising direction of research. Translational neurodegeneration. 2012;1(1):11.

22. Ross GW, Petrovitch H, Abbott RD, Tanner CM, Popper J, Masaki K, et al. Association of olfactory dysfunction with risk for future Parkinson’s disease. Annals of neurology. 2008;63(2):167–173.

23. Haehner A, Hummel T, Hummel C, Sommer U, Junghanns S, Reichmann H. Olfactory loss may be a first sign of idiopathic Parkinson’s disease. Movement disorders. 2007;22(6):839–842.

24. Postuma RB, Lang AE, Massicotte-Marquez J, Montplaisir J. Potential early markers of Parkinson disease in idiopathic REM sleep behavior disorder. Neurology. 2006;66(6):845–851.

25. Wetzel PA, Baron MS, Gitchel GT. Automated analysis system for the detection and screening of neurological disorders and deficits; 2014. Available from: https://www.google.com/patents/WO2014159498A3?cl=en.

26. Hufschmidt HJ. Proprioceptive origin of parkinsonian tremor. Nature. 1963;.

27. Stiles RN, Pozos RS. A mechanical-reflex oscillator hypothesis for parkinsonian hand tremor. Journal of applied physiology. 1976;40(6):990–998.

28. Stein R, Oĝuztöreli M. Tremor and other oscillations in neuromuscular systems. Biological cybernetics. 1976;22(3):147–157.

29. Rack P, Ross H. The role of reflexes in the resting tremor of Parkinson’s disease. Brain. 1986;109(1):115–141.

30. Schnider S, Kwong R, Lenz F, Kwan H. Detection of feedback in the central nervous system using system identification techniques. Biological cybernetics. 1989;60(3):203–212.

31. Beuter A, Milton J, Labrie C, Glass L, Gauthier S. Delayed visual feedback and movement control in Parkinson’s disease. Experimental neurology. 1990;110(2):228– 235.

32. Beuter A, Bélair J, Labrie C. Feedback and delays in neurological diseases: a modeling study using dynamical systems. Bulletin of mathematical biology. 1993;55(3):525–541.

33. Deuschl G, Raethjen J, Baron R, Lindemann M, Wilms H, Krack P. The pathophysiology of parkinsonian tremor: a review. Journal of neurology. 2000;247(5):V33– V48.

34. Rodriguez-Oroz MC, Jahanshahi M, Krack P, Litvan I, Macias R, Bezard E, et al. Initial clinical manifestations of Parkinson’s disease: features and pathophysiological mechanisms. The Lancet Neurology. 2009;8(12):1128–1139.

35. Zhang D, Poignet P, Bo AP, Ang WT. Exploring peripheral mechanism of tremor on neuromusculoskeletal model: A general simulation study. IEEE Transactions on Biomedical Engineering. 2009;56(10):2359–2369.

36. Hallett M. Tremor: pathophysiology. Parkinsonism & related disorders. 2014;20:S118–S122.

37. Palanthandalam-Madapusi HJ, Goyal S. Is Parkinsonian Tremor a limit cycle? Journal of Mechanics in Medicine and Biology. 2011;11(05):1017–1023.

38. Milton JG. Time delays and the control of biological systems: An overview. IFAC-PapersOnLine. 2015;48(12):87–92.

39. Shah VV, Goyal S, Palanthandalam-Madapusi HJ. Clinical Facts Along With a Feedback Control Perspective Suggest That Increased Response Time Might Be the Cause of Parkinsonian Rest Tremor. Journal of Computational and Nonlinear Dynamics. 2017;12(1):011007.

40. Chumacero E, Yang J, Chagdes JR. Effect of sensory-motor latencies and active muscular stiffness on stability for an ankle-hip model of balance on a balance board. Journal of biomechanics. 2018;75:77–88.

41. Edwards R, Beuter A, Glass L. Parkinsonian tremor and simplification in network dynamics. Bulletin of mathematical biology. 1999;61(1):157–177.

42. Elble RJ. Central mechanisms of tremor. Journal of clinical neurophysiology. 1996;13(2):133–144.

43. Sak W. The Croonian Lectures on Some Disorders of Motility and of Muscle Tone, With Special Reference to the Corpus Striatum. The Lancet. 1925;206(5314):1 – 10.

44. Evarts E, Teräväinen H, Calne D. Reaction Time in Parkinson’s Disease. Brain: A Journal of Neurology. 1981;104(1):167–186.

45. Bloxham C, Dick D, Moore M. Reaction Times and Attention in Parkinson’s Disease. Journal of Neurology, Neurosurgery & Psychiatry. 1987;50(9):1178–1183.

46. Heilman KM, Bowers D, Watson RT, Greer M. Reaction Times in Parkinson Disease. Archives of Neurology. 1976;33(2):139–140.

47. Shah VV, Goyal S, Palanthandalam-Madapusi HJ. A Possible Explanation of How High-Frequency Deep Brain Stimulation Suppresses Low-Frequency Tremors in Parkinson’s Disease. IEEE Transactions on Neural Systems and Rehabilitation Engineering. 2017;25(12):2498–2508.

48. Sarkar TK, Pereira O. Using the matrix pencil method to estimate the parameters of a sum of complex exponentials. Antennas and Propagation Magazine, IEEE. 1995;37(1):48–55.

49. Kourou K, Exarchos TP, Exarchos KP, Karamouzis MV, Fotiadis DI. Machine learning applications in cancer prognosis and prediction. Computational and structural biotechnology journal. 2015;13:8–17.

50. Cortes C, Vapnik V. Support-vector networks. Machine learning. 1995;20(3):273– 297.

51. Tang J, Alelyani S, Liu H. Feature selection for classification: A review. Data Classification: Algorithms and Applications. 2014; p. 37.

52. Kristiaan P, Hong-Li Z. Support Vector Machines for Longitudinal Analysis. 2015;.

53. De Lima ER, Andrade AO, Pons JL, Kyberd P, Nasuto SJ. Empirical mode decomposition: a novel technique for the study of tremor time series. Medical and biological engineering and computing. 2006;44(7):569–582.

54. Gitchel GT, Wetzel PA, Baron MS. Pervasive ocular tremor in patients with Parkinson disease. Archives of neurology. 2012;69(8):1011–1017.

55. Flowers K, Downing A. Predictive control of eye movements in Parkinson disease. Annals of neurology. 1978;4(1):63–66.

56. Senders JW, Cruzen M. Tracking performance on combined compensatory and pursuit tasks. Wright Air Development Center, Air Research and Development Command, United States Air Force; 1952.

57. Giza E, Fotiou D, Bostantjopoulou S, Katsarou Z, Karlovasitou A. Pupil light reflex in Parkinson’s disease: evaluation with pupillometry. International Journal of Neuroscience. 2011;121(1):37–43.

58. Micieli G, Tassorelli C, Martignoni E, Pacchetti C, Bruggi P, Magri M, et al. Disordered pupil reactivity in Parkinson’s disease. Clinical Autonomic Research. 1991;1(1):55–58.

59. You S, Hong JH, Yoo J. Analysis of pupillometer results according to disease stage in patients with Parkinson’s disease. Scientific reports. 2021;11(1):1–6.

60. Heenan M, Scheidt RA, Woo D, Beardsley SA. Intention tremor and deficits of sensory feedback control in multiple sclerosis: a pilot study. Journal of neuroengineering and rehabilitation. 2014;11(1):170.

61. Thompson JD, Franz JR. Do kinematic metrics of walking balance adapt to perturbed optical flow? Human Movement Science. 2017;54:34–40.

62. Franz JR, Francis CA, Allen MS, O’Connor SM, Thelen DG. Advanced age brings a greater reliance on visual feedback to maintain balance during walking. Human movement science. 2015;40:381–392.

63. Cona F, Pizza F, Provini F, Magosso E. An improved algorithm for the automatic detection and characterization of slow eye movements. Medical engineering & physics. 2014;36(7):954–961.

64. King RC, Villeneuve E, White RJ, Sherratt RS, Holderbaum W, Harwin WS. Application of data fusion techniques and technologies for wearable health monitoring. Medical Engineering & Physics. 2017;.

65. Meel-van den Abeelen AS, van Beek AH, Slump CH, Panerai RB, Claassen JA. Transfer function analysis for the assessment of cerebral autoregulation using spontaneous oscillations in blood pressure and cerebral blood flow. Medical engineering & physics. 2014;36(5):563–575.

66. Panerai R, Rennie J, Kelsall A, Evans D. Frequency-domain analysis of cerebral autoregulation from spontaneous fluctuations in arterial blood pressure. Medical and Biological Engineering and Computing. 1998;36(3):315–322.

